# Whole genome sequence of an edible and potential medicinal fungus, *Cordyceps guangdongensis*

**DOI:** 10.1101/254243

**Authors:** Chenghua Zhang, Wangqiu Deng, Wenjuan Yan, Taihui Li

## Abstract

*Cordyceps guangdongensis* is an edible fungus which has been approved as a Novel Food by the Chinese Ministry of Public Health in 2013. It also has a broad application prospect in pharmaceutical industries with many medicinal activities. In this study, the whole genome of *C. guangdongensis* GD15, a single spore isolate from a wild strain, was sequenced and assembled with Illumina and PacBio sequencing technology. The generated genome is 29.05 Mb in size, comprising 9 scaffolds with an average GC content of 57.01%. It is predicted to contain a total of 9150 protein-coding genes. Sequence identification and comparative analysis indicated that the assembled scaffolds contained two complete chromosomes and four single-end chromosomes, showing a high level assembly. Gene annotation revealed a diversity of transporters that could contribute to the genome size and evolution. Besides, approximately 15.49% and 13.70% genes involved in metabolic processes were annotated by KEGG and COG respectively. Genes belonging to CAZymes accounted for a proportion of 2.84% of the total genes. In addition, 435 transcription factors (TFs) were identified, which were involved in various biological processes. Among the identified TFs, the fungal transcription regulatory proteins (18.39%) and fungal-specific TFs (19.77%) represented the two largest classes of TFs. These data provided a much needed genomic resource for studying *C. guangdongensis*, laying a solid foundation for further genetic and biological studies, especially for elucidating the genome evolution and exploring the regulatory mechanism of fruiting body development.

## INTRODUCTION

*Cordyceps guangdongensis* T. H. Li, Q. Y. Lin & B. Song (*Cordycipitaceae*) was discovered in southern China (Lin *et al*. 2008), and has been successfully cultivated (Lin *et al*. 2010). Its fruiting body is nontoxic, and has become the second Novel Food of *Cordyceps* species approved by the Ministry of Public Health of China in 2013. This fungus has rich nutrient contents and bioactive compounds, such as cordycepic acid, adenosine and polysaccharide (Lin *et al*. 2009; 2010). The contents of the above-mentioned bioactive compounds in *C. guangdongensis* are similar to those of the traditional Chinese invigorant, *Ophiocordyceps sinensis* (=*Cordyceps sinensis*) (Lin *et al*. 2009). Previous researches by the authors’ group indicated that the fruiting bodies of *C. guangdongensis* possessed various therapeutic properties, including antioxidant activity (Zeng *et al*. 2009), longevity-increasing activity (Yan *et al*. 2011), anti-fatigue effect (Yan *et al*. 2013), curative effect on chronic renal failure (Yan *et al*. 2012), and anti-inflammatory effect (Yan *et al*. 2014). These active effects provided great potential for its application in food and medicinal industries. However, it is worth considering what makes the fruiting bodies of *C. guangdongensis* so valuable and how to improve the fruiting body yield. Therefore, it is a matter of cardinal significance for further understanding the fruiting body development and metabolism mechanisms of *C. guangdongensis*, as well as the evolutionary relationship between other related species.

In recent years, whole genome sequencing (WGS) is widely used to analyze the relevance of phenotypic characters and genetic mechanism. The advanced sequencing techniques and rapidly developed bioinformatic methods had become the practical ways to further explore the molecular mechanisms of fungal development and metabolism, the systematic taxonomy and the evolutionary relationship of fungi. To date, numerous genomes of fungi under Hypocreales have been released in Ensembl fungus database (http://fungi.ensembl.org/index.html). Based on the available genome sequences, researchers not only ascertained evolutionary relationship of many fungi (Zheng *et al*. 2011; Bushley *et al*. 2013; Xia *et al*. 2017), but also identified quite a lot medically active components (Zheng *et al*. 2011; Yin *et al*. 2012; Bushley *et al*. 2013; Quandt *et al*. 2015). Meanwhile, various transcription factors (TFs), including bZIP TFs, zinc finger TFs and fungal-specific TFs, were proved to be involved in fruiting body development by transcriptome analysis on the basis of genome sequences (Zheng *et al*. 2011; Yin *et al*. 2012; Yang *et al*. 2016). However, the whole genome sequence for *C. guangdongensis* is still lacking.

In order to acquire abundant molecular information for further effectively exploring the relevance of phenotypic characters and genetic mechanisms in *C. guangdongensis*, the whole genome of *C. guangdongensis* was firstly sequenced by a combined Illumina HiSeq2000 and PacBio platform in this study. The assemble level, the types of transposable elements and the transcriptional factors (TFs) were further analyzed. This genomic resource provided a novel insight into further genetic researches that will facilitate in its application addressed in many areas of biology.

## MATERIAL AND METHODS

### Fungal strains and DNA extraction

Sample used for whole genome sequencing and assembly was isolated from the strain GDGM30035 (wild fruiting bodies of *C. guangdongensis*). The strain was cultured on the PDA medium at 23±1° for four weeks, aqueous suspensions of fungal spores were prepared by pouring sterile distilled water onto the sporulated cultures and gently scrapping the agar surface. The spore suspension was collected by passing through four layers of sterile cheesecloth to remove mycelial fragments. Spore suspension was adjusted to 1×10^3^ conidia ml/l using a hemocytometer, and was coated on the PDA medium covered with cellophane. Single colony (*C. guangdongensis* GD15) was transferred onto new PDA medium and subcultured for three times. For DNA isolation, the strain was cultured on PDA medium, which was then covered with cellophane at 23°. Genomic DNA from 7-day-old fungal colony was extracted using a CTAB-based extraction buffer (Watanabe *et al*. 2010). The DNA concentration was determined using UV-Vis spectrophotometer (BioSpec-nano), and the integrity of the DNA was detected using a 0.8% agarose gel, while the purity of the DNA was analyzed by a PCR program using 16S rDNA primers.

### Genome sequencing and assembly

The genome of *C. guangdongensis* GD15 was sequenced with the hybrid of the next generation sequencing technology Illumina HiSeq 2000 and the third-generation sequencing platform, PacBio, at the Beijing Genomics Institute at Shenzhen. DNA libraries with 500 bp inserts were constructed and sequenced with the Illumina HiSeq2000 Genome Analyzer sequencing technique. Long insert SMRTbell template librarieswerepreparedaccordingtoPacBioprotocols.Theunqualifiedrawreads obtained by PacBio were filtered out, the subreads (≥1000bp) were corrected by Proovread 2.12 (https://github.com/BioInf-Wuerzburg/proovread) and then were initially assembled by SMRT Analysis v.2.3.0 (Chin *et al*. 2013). The preliminary assembly results were further corrected using small Illumina reads by GATK v1.6-13 (http://www.broadinstitute.org/gatk/), the scaffolds were assembled and optimized using long Illumina reads by SSPACE Basic v2.0 (http://www.baseclear.com/genomics/bioinformatics/basetools/SSPACE) and PBJelly2 v15.8.24 (https://sourceforge.net/projects/pb-jelly).

### Genome components analysis

The characteristic telomeric repeats (TTAGGG/CCCTAA) were searched on the both ends of each scaffold within 100bp length. Repetitive elements included tandem repeats and transposable elements (TEs). Tandem repeats were searched in all scaffolds by Tandem Repeats Finder (TRF 4.04) described by Benson (1999). TEs annotation was performed by RepeatMasker 4.06 (Smit *et al*. 2014) based on the Repbase database (http://www.girinst.org/repbase). The tRNAs were predicted by tRNAscan-SE 1.3.1 (Lowe and Eddy, 1997), rRNAs were identified using RNAmmer 1.2 (Lagesen *et al*. 2007), and sRNA were predicted by Infernal based on the Rfam database (Gardner *et al*. 2009). Genes were annotated based on sequence homology and *de novo* gene predictions. The homology approach was based on the reference genomes download from EnsemblFungi (http://fungi.ensembl.org/index.html) including the protein sequences of *C. militaris*, *O. sinensis* and *Cordyceps ophioglossoides* (=*Tolypocladium ophioglossoides*). The *de novo*gene predictions were performed by Genemarkes 4.21 (Ter-Hovhannisyan *et al*. 2008).

### Functional annotation

Structural and functional annotations of genes were performed according to various databases of ARDB (Antibiotic Resistance Genes Database) (Liu *et al*. 2009), CAZymes (Carbohydrate-Active enZYmes Database) (Cantarel *et al*. 2009), COG (Cluster of Orthologous Groups) (Tatusov *et al*. 2003), GO (Gene Ontology) (Bard and Winter 2000), KEGG (Kyoto Encyclopedia of Genes and Genomes) (Kanehisa *et al*. 2006), NR (Non-Redundant Protein Database), P450 (Magrane *et al*. 2011), PHI (Pathogen Host Interactions) (Torto-Alalibo *et al*. 2009), SwissProt (Magrane *et al*. 2011), T3SS (Type III secretion system Effector protein) (Vargas *et al*. 2012), TrEMBL, VFDB (Virulence Factors of Pathogenic Bacteria) (Chen *et al*. 2016)and so on. Transcription factors (TFs) were analyzed based on the TRANSFAC database (http://www.gene-regulation.com/pub/databases.html#transfac).

### Data availability

The genome sequencing project has been deposited at GenBank under the accession number NRQP00000000. The BioProject designation for this project is PRJNA399600.

## RESULTS AND DISCUSSION

### Whole-genome assembly

A total of 3,926,378,523 reads representing a cumulative size of 3.926 Gb were generated, including 13,392,532 and 3,912,985,991 reads obtained from Illumina and PacBio sequencing platforms, respectively. The PacBio sequencing results showed high quality of polymerasereads and subreads (Fig.S1). After the low quality reads were filtered, a total of 3,484,503,143 reads were assembled into 9 scaffolds with an N50 of 7.88 Mb from ∼183 average coverage, and a total length of 29.05 Mb with a 57.01% GC content was obtained (Fig. 1A). Based on the Illumina sequencing data, the predicted genome size by K-mer analysis was 31.58Mb, the total size of the combined assembly closely matched this estimate size (91.98%). Besides, compared to the previously reported draft genomes listed in Table 1, the new assembly sequences were more contiguous with higher GC content.

**Figure 1.**
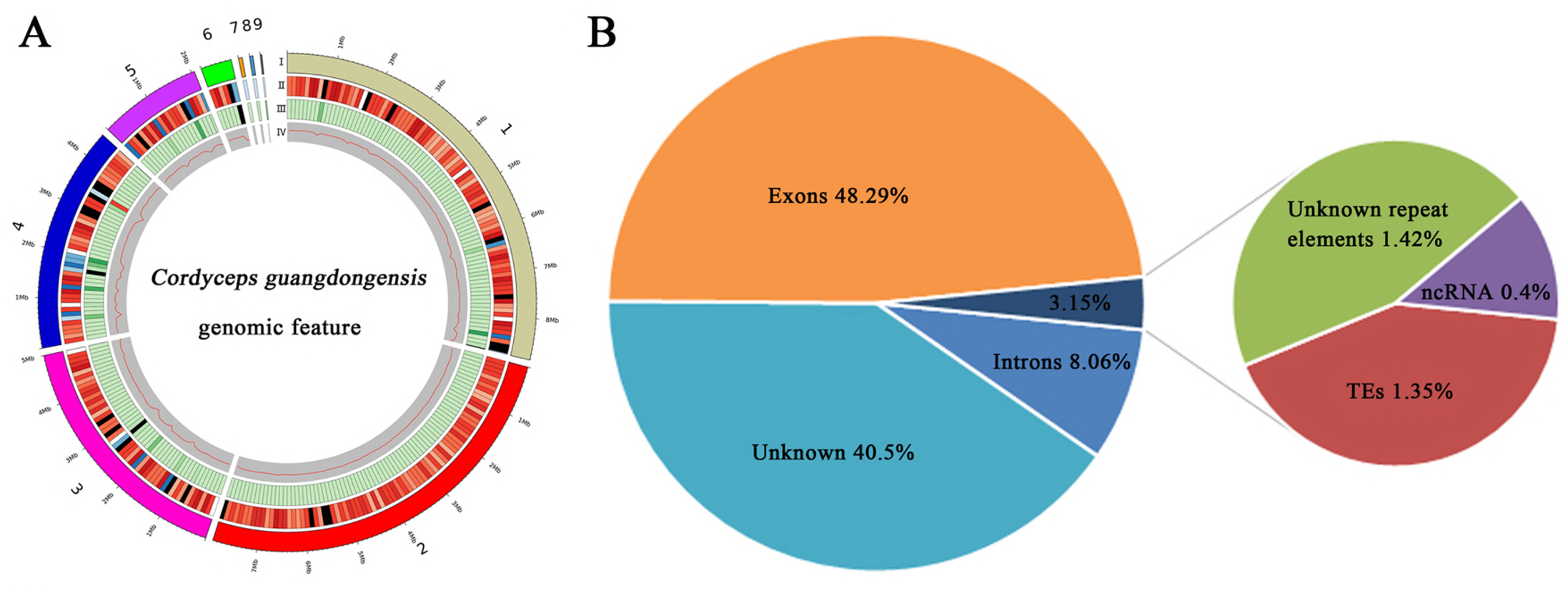
General genomic feature of *Cordyceps guangdongensis*. A, I, scaffolds; II, gene density represented as the number of genes per 100 kb; III, percentage of coverage of repetitive sequences; I V, GC content estimated by the percentage of G + C in 100 kb. B, Genomic element density including genic and nongenic features of the overall genome assembly length including 40.5% non-annotated sequences.

### Chromosome analysis

Sequences analysis of telomeric repeats was used to estimate the number of chromosomes for *C. guangdongensis* genome according to the method described by Zheng *et al*. (2011). The characteristic telomeric repeats (TTAGGG/CCCTAA)_n_ were found at either 5’ or 3’ terminal of 6 scaffolds, of which the telomeric repeats were found at both ends of scaffolds 1 and 3, suggesting that the two scaffolds are complete chromosomes. The lengths of the two complete chromosomes were about 8.81M and 5.00M, respectively. Single-ended telomeric repeats were found at four scaffolds, including the start of scaffolds 4 and 6, the end of scaffolds 2 and 5, suggesting that these four scaffolds extended to the telomeres. The lengths of the four candidate scaffolds are about 7.88M, 4.50M, 2.05M and 0.61M, respectively. The remaining three scaffolds contained none of telomeric repeats, possibly due to incompleteness of the scaffold sequence data (Table 2, Table S1). Furthermore, on scaffold 7, 14 genes were identified which belonged to the core genes of mitochondrial genome, indicating that this scaffold represent mitochondrial genome sequence. Previous researches showed that the haploid genome of *C. militaris* contains seven chromosomes (Kramer *et al*. 2017), and *Cordyceps subsessilis* (=*Tolypocladium inflatum*) also contains seven chromosomes (Stimberg *et al*. 1992). Taking into consideration of the chromosome number in the above related species and the present telomeric repeats analysis, it was inferred that *C. guangdongensis* may possibly also contain seven chromosomes, and this hypothesis should be further proved by karyotype analysis.

### Genome features and annotation

As shown in Table 3, a total of 9150 protein-coding genes were predicted in the genome, including 31 rRNA, 111 tRNA, 121 sRNA, 25 snRNA and 26 miRNA. The cumulative length of total genes accounted for 56.35% of the whole genome sequence length, and the lengths of most genes were in the range of 200-5000bp (Fig. S3). The exons showed a large proportion (48.29%) with the max number of 29,548, and the number of introns was 20,398, with a total length of 2.34M (8.06%). The ncRNA with a total number of 314 represented 0.4% of the genome assembly, suggesting that they contributed only a small proportion to the overall genome size (Fig. 1B).

Of the 9150 identified genes, 8486 genes (92.74%) were annotated by the databases described in the methods (Table S2). The present paper was mainly focused on the genes involved in metabolic process. Among all the predicted genes, approximately 37.22% were annotated by KEGG pathway, including 15.49% involved in metabolism, accounting for the major proportion. Genes classified into functional categories based on the COG analysis accounted for 23.83%, including 13.70% involved in metabolic processes. Thereinto, approximately 2.01% of total genes were related to the biosynthesis, transportation, and catabolism of secondary metabolites (Fig. 2). The percentage of genes encoding CAZymes was 2.84%, which contributed to substrate degradation processes as nutrition for fungal development and reproduction. Among the genes related with CAZymes, 103 genes encoding glycoside hydrolases (GHs) accounted for the largest proportion of 1.12% of the total genes, followed by 78 genes encoding carbohydrate-binding modules (CBMs) accounting for 0.85%,.and then 66 genes encoding glycosyl transferases (GTs) with a percentage of 0.72%. Genes belong to the carbohydrate esterases (CEs) and polysaccharide lyases (PLs) had much lower percentages of about 0.13% and 0.01%, respectively. Since the genes relevant to CAZymes in *Pleurotus eryngii* were not only involved in XXXX, but also involved in its primordium differentiation and fruiting body development (Xie *et al*. 2017), the above identified genes of CAZymes in *C. guangdongensis* could likely also be involved in the primordium differentiation and fruiting body development, and the studied results in this paper will be beneficial to further study on the genetic and molecular mechanisms of the fruiting body development.

**Figure 2.**
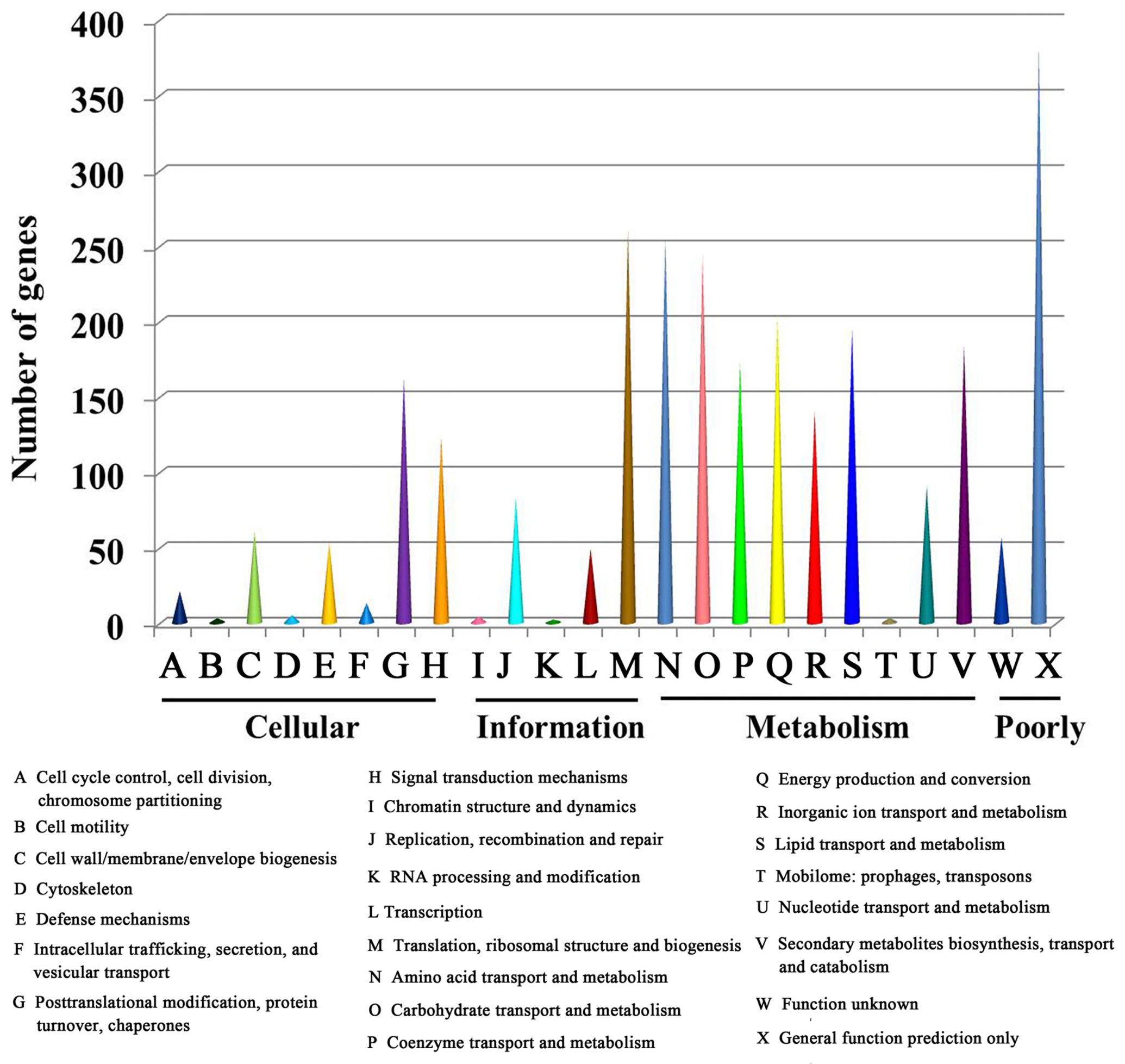
COG functional classification of proteins in the *Cordyceps guangdongensis* genome.

### Repetitive elements

The cumulative sequences of repetitive elements identified in *C. guangdongensis* genome occupied a percentage of 2.77% of the assembly sequences. The tandem repeats represented 1.42% of the genome assembly with a total length of 412,989 bp; and the TEs represented 1.35% of the genome assembly with a total length of 393,608 bp. The total number of TE families analyzed by RepeatMasker in the genome assembly is 1534, of which 1527 (99.5%) belonged to the known TEs, including 1033 retrotransposons (Class I) and 494 DNA transposons (Class II); while the remaining TEs could not be classified at this moment (Table 4 and Table S3).

The retrotransposons in Class I can be mainly divided into three groups of TEs, including LINE, LTR and SINE; and each group contains several subgroups. Retrotransposons, particularly L1, Copia, DIRS, ERV1, Gypsy, Pao, Alu and so on, are the easiest ones to be annotated, which are also the most abundant transposons in fungi (Suarez *et al*. 2017). The DNA transposons in Class II contain a lot of known groups and some unclassified members. Among the transposons, hAT, MULE, PIF-Harb and Tc1-Mariner were reported to be extraordinarily abundant in fungi, while the transposons CMC and piggyBac have limited taxonomic distribution and seem to remain in a few fungal taxa (Muszewska *et al*. 2017). Other transposons, including P, Sola, Dada, Ginger, Zisupton and Merlin, had been identified only in a handful of species (Kojima and Jurka 2013; Iyer *et al*. 2014; Majorek *et al*. 2014).

Previous studies indicated that TEs contributed to genome size expansion and evolution (Cordaux *et al*. 2009; Sun *et al*. 2012), and played a crucial roles in a wide range of biological events, including organism development (Kano *et al*. 2009; Garcia-Perez *et al*. 2016), regulation (Elbarbary *et al*. 2016) and differentiation (Morales-Hernandez *et al*. 2016), sometimes acting as novel promoters to activate transcriptive process (Faulkner *et al*. 2009; Mita *et al*. 2016). Therefore, the abundance in *C. guangdongensis* would have more significance in this regard, and they should be further noticed for studying both fungal taxonomy application and regulatory roles in fruiting body development by bonding to the transcription factors.

### Transcription factors

Transcription factors (TFs) are essential to modulating diverse biological processes by regulating gene expression and play central roles in the development and evolution. In this study, function annotation identified 435 genes to be TFs in *C. guangdongensis*, accounting for 4.75% of the total predicted genes. Like other fungi, genes encoding fungal-specific TFs (86 members) and fungal transcription regulatory proteins (80 members) represented the two largest classes of TFs in *C. guangdongensis*, followed by C_2_H_2_-type Zinc finger TFs (54 members) and Winged helix-repressor DNA binding proteins (54 members), accounting for approximately 19.77%, 18.39%, 12.41% and 12.41% of the total predicted TFs, respectively (Fig. 3).

**Figure 3.**
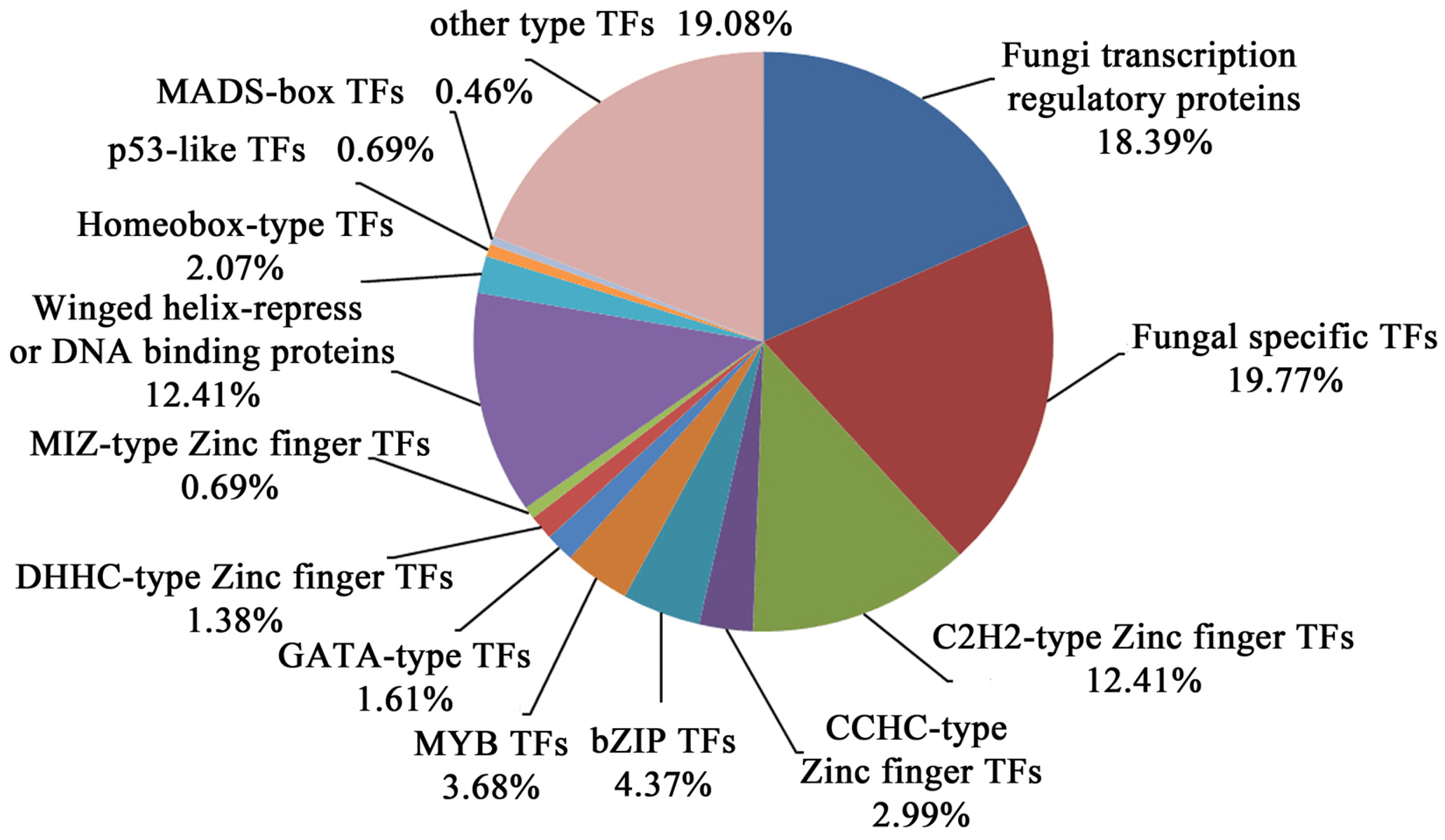
Transcription factors analysis in the *Cordyceps guangdongensis* genome.

Moreover, other different types of Zinc finger transcription factors were identified, including 13 CCHC-type Zinc finger TFs (2.98%), 6 DHHC-type TFs (0.06%) and 3 MIZ-type TFs (0.03%). Apart from these, there were 19 bZIP TFs (0.21%), 16 MYB TFs (0.17%), 7 GATA TFs (0.08%) and 9 Homeobox-type TFs (0.10%). In *C. militaris*, majority of TFs, such as Zn_2_Cys6-type TFs, GATA-type TFs, bZIP TFs, CHCC-type TFs, were differential expressions during fruiting body developmental stages (Zheng *et al*. 2011). Hence, the known information about various TFs could guide the search for exploring the regulatory mechanism of TFs on fruiting body development in *C. guangdongensis*.

## Conclusion

High quality of genome sequencing of *C. guangdongensis* was presented in this study. Two complete chromosomes and four single-end chromosomes were assembled through sequence analysis of the genome. In the genomic sequences, diversities of transposable elements were identified, which may contribute to the genome size and evolution. Moreover, transcription factors existing in *C. guangdongensis* were identified and classified, which may facilitate the further studying of fruiting body development. With the knowledge of the whole genome sequence, more detailed molecular information will be revealed, providing interesting insights into better understanding the relevance of phenotypic characters and genetic mechanism in *C. guangdongensis*.

## ACKNOWLEDGMENTS

This work was supported by GDAS’ Special Project of Science and Technology Development (No. 2017GDASCX-0822), Natural Science Foundation of Guangdong Province, China (2017A030310533), the Science and Technology Planning Project of Guangdong Province, China (No. 2015A030302052; 2016A030303035), the National Natural Science Foundation of China (No. 31470155), and the Science and Technology Planning Project of Guangzhou, China (No. 201504291620324). The authors sincerely thank the professor Chengshu Wang in Shanghai Institute for Biological Sciences for providing the guidance of genome sequencing.

---

**Table 1** Assembly summary statistics of *Cordyceps guangdongensis* GD15 compared to other *Cordyceps* genomes.

**Table 2**Chromosome analysis of *Cordyceps guangdongensis* GD15 genomic sequence.

**Table 3** Genome annotation features of *Cordyceps guangdongensis* GD15.

**Table 4**Class analysis of transposable elements repeats in *Cordyceps guangdongensis*.

**Table S1** Genome sequences used to analyze the chromosome

**Table S2** Genome annotation of proteins in *Cordyceps guangdongensis* genome

**Table S3** Transposable elements classification in *Cordyceps guangdongensis*

## Figure legends

**Figure S1** Length and quality distributions of PacBio reads.

**Figure S2** Length statistics of scaffolds generated by genomic sequencing.

**Figure S3**Distribution of genelength predicted in the*Cordyceps guangdongensis* genome.

